# HEAD AND SHOULDERS - THE IMPACT OF AN EXTENDED HEAD MODEL ON THE SIMULATION AND OPTIMIZATION OF TRANSCRANIAL ELECTRIC STIMULATION

**DOI:** 10.1101/2024.08.29.610251

**Authors:** Sybren Van Hoornweder, Vittoria Cappozzo, Laura De Herde, Oula Puonti, Hartwig R. Siebner, Raf L.J. Meesen, Axel Thielscher

## Abstract

Electric field calculations are increasingly used for dose characterization of transcranial electrical stimulation (tES), but existing open-source head models are inaccurate for extracephalic montages that include electrodes placed on the neck or shoulder. We introduce the “Ernie Extended” model, an MRI- and CT-derived open-source head model extending to the upper shoulder region. Simulations of extracephalic tES targeting the cerebellum and supplementary motor area show significant differences in electric fields when using Ernie Extended compared to the non-extended Ernie model. Additionally, we propose an electrode layout that complements the electroencephalography 10– 20 system with extracephalic electrode positions. We demonstrate the use of this layout for optimizing multi-electrode tES montages for cerebellar stimulation, enhancing focality and reducing off-target stimulation, particularly of the spinal cord. Our results highlight the practical value of the Ernie Extended model for accurately characterizing doses produced by extracephalic tES montages and when targeting more caudal brain regions.

## 1. Introduction

Electric field simulations are essential for the characterization and optimization of the current flow induced by transcranial electric stimulation (tES) [1-5]. Existing open-source simulation tools are based on volume conductor models that cover solely the head. Yet, tES montages may extend to regions outside the head, such as the neck or upper shoulders, for instance when tES is used to target the cerebellum [6-8]. In these cases, electric field simulations currently have to rely on approximations such as placing the “extracephalic” electrode at the bottom of the volume conductor model as close as possible to its actual location (e.g., [9]). However, the accuracy of these approximations remains unclear, and previous work has demonstrated the importance of precise anatomical and montage representations for current flow modeling [10, 11]. Thus, there is a need for extended head models. Although these have been previously developed [11-16], none are open-source and many face shortcomings that impact current flow modeling accuracy, such as a lack of grey matter gyrification or the reliance on solely T1w MRI data (cf., Appendix 1 for a comparison) [3, 10, 17, 18].

We introduce the “Ernie Extended” model, an open-source, anatomically accurate, extended head model. Ernie Extended is derived from T1w-, T2w-MRI and CT data, and includes 13 tissue types. We highlight the value of the new model by comparing extracephalic tES electrical field simulations based on Ernie Extended with simulations based on the non-extended Ernie model. Also, we explore leadfield-based optimization for cerebellar tES based on an extended electrode layout including extracephalic positions. The Ernie Extended model and an accompanying tutorial based on this work will be made freely available through the SimNIBS website.

## 2. Methods

### 2.1. Data acquisition

We acquired high-resolution fat-suppressed T1w- and T2w-MRI scans and a CT scan of the head, along with a non-fat-suppressed T1w MRI scan of the head and shoulders for a single individual (cf., Appendix 2). All scans were co-registered to the T1w head scan using FLIRT and elastix for, respectively, linear and non-linear registration [19-22]. Non-linear registration was needed to align the neck between the MRI and the CT scans as participant placement in the scanners differed.

The study was approved by the Ethical Committee of the Capital Region of Denmark, and written informed consent was obtained from all participants prior to the scans. Moreover, the participant had no previous history of neurological or psychiatric disorders and was screened for contraindications to MRI and CT.

### 2.2. Extended Head Segmentation, Processing and Meshing

The segmentation was performed for three regions separately: upper head with skull, lower head and neck, and the shoulder region.

For the upper head region, the T1w- and T2w-MRI head scans and CT-scan were segmented via SimNIBS-CHARM [23, 24]. The compact bone of the MRI-based segmentation was replaced by that of the CT-based segmentation. Remaining non-assigned voxels in the resulting segmentation were filled using neighboring voxel labels. Similarly, the brain representation was improved by using the pial and white matter surfaces reconstructed by FreeSurfer 7.3 from the high-resolution T1w-scan [25].

For the lower region, including the neck, we used the T1w- and T2w-MRI head scan-based SimNIBS-CHARM segmentations, the CT-scan, and the T1w head and shoulders scan. Semi-automated segmentation started from the MRI-segmentation of the head, with the cervical vertebrae (compact and spongy bone) and the esophagus (air) being based on thresholded CT data. The intervertebral discs were segmented manually. Fat and non-fat tissue (mostly muscle) were distinguished by thresholding the non-fat-suppressed T1w-image of the head and shoulders. Visual inspection and manual correction of all tissues was done using ITK-SNAP [26] and FreeView [25]. The same programs were used for the manual segmentation of the shoulder region.

The three segmentations, consisting of 13 tissues, were combined and morphological operations and Gaussian smoothing were used with tissue-specific parameters for the head and shoulders. For the head and neck, post-processing was minimal. For the shoulders, more post-processing was applied to mitigate staircasing due to manual segmentation, albeit we also aimed to minimize smoothing to avoid the loss of anatomical details. The final segmentation was meshed into a tetrahedral mesh using the SimNIBS meshmesh command (**Figure 1A**).

**Figure 1.**
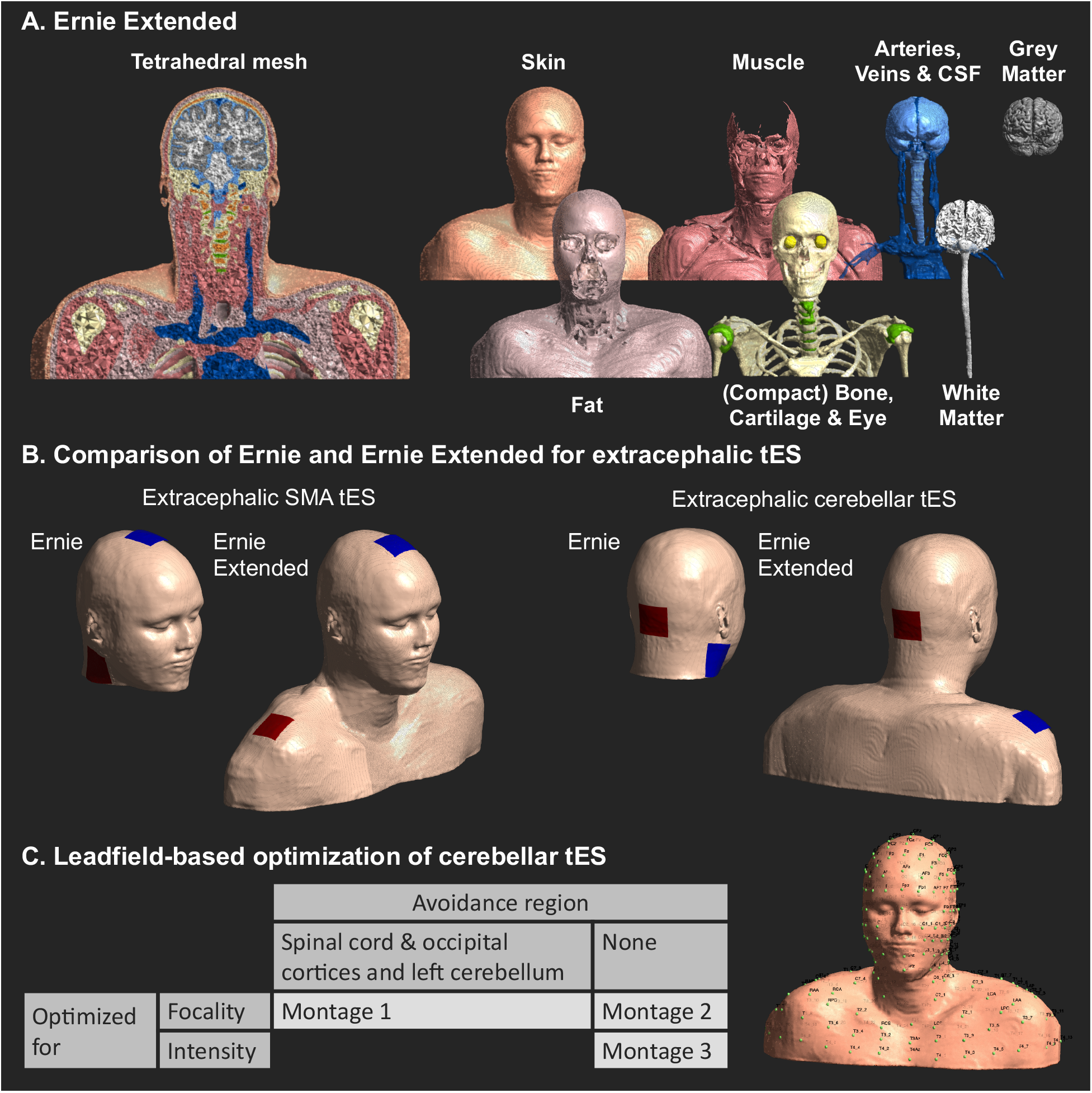
The panels show Ernie Extended, the simulated tES montages, and the performed cerebellar tES optimizations. **A)** The Ernie Extended mesh **B)** tES montages used in Ernie Extended (upper row) and Ernie (lower row). **C)** Three leadfield-based optimizations for cerebellar tES were done in Ernie Extended.

### 2.4. Effect of Extended Head Models on tES electric field simulations

We tested whether approximations of two extracephalic tES montages using a non-extended head model, Ernie, yield similar electric fields as simulations with Ernie Extended. Other than stopping at the chin, Ernie is identical to Ernie Extended (**Figure 1B**). Standard conductivity values were used for all tissues (Appendix 3).

The first montage, used for cerebellar tES [6-8], places the anode ∼2 cm below the inion, and the cathode over the right shoulder. The second montage, used in persons with obsessive compulsive disorder [27], positions the cathode over the supplementary motor area (SMA) and the anode over the left shoulder. For consistency, the right shoulder was used in both montages with a stimulation intensity of 1 mA. We warped all electric field distributions to MNI space, and calculated voxel-wise difference maps between the extended and non-extended models. We also extracted the 99.9^th^ percentile electric field magnitude in grey matter [28].

### 2.5. Effect of Extended Head Models on tES Optimization

Ernie Extended expands electrode locations for leadfield-based tES optimization– a method to determine the optimal tES electrode placement for a given target [29, 30]– to extracephalic locations. We complemented the EEG 10-20 layout by adding evenly distributed positions in transversal planes defined per vertebral level, and several palpable bony landmarks such as the coracoid process (**Figure 1C**).

Then, we explored the benefits of the extended electrode layout for optimizing multi-electrode montages targeting the cerebellum, which is challenging with standard head models stopping at the chin. Following leadfield calculations, three optimizations were performed for the right cerebellum (coordinate: 17.74, -54.46, -32.00, 5 mm radius): montage 1 and 2 were optimized for focality, aiming to induce electric field magnitude = 0.2 V/m in the target, and montage 3 was optimized for maximizing the field magnitude in the target. Montage 1 avoided electric fields in the spinal cord and right occipital cortex (**Figure 1C**). Maximum current intensity was 4 mA, with a maximum of 2 mA per electrode and maximally 8 active electrodes. Per montage, we used spherical regions of interest to extract mean electric field magnitude in the right and left cerebellum, the upper spinal cord and right occipital cortex regions.

## 3. Results

### 3.1. Differences of the electric field distributions in the extended and non-extended models

Extracephalic tES montages targeting the cerebellum and SMA were simulated using Ernie and Ernie Extended (**Figures 2A** and **2B)**. While peak magnitudes were similar in Ernie (0.242 V/m) and Ernie Extended (0.253 V/m) for cerebellar tES, the area exceeding the 99.9^th^ percentile electric field magnitude differed, indicating spatial differences in the electric field distribution. In Ernie Extended, the field peaks occurred in the caudal cerebellum, while in Ernie, the fields were strongest in the right, rostral, latero-inferior temporal cortex. Overall, the simulated electric field magnitude was stronger in the right temporal cortex when using the Ernie model, while the Ernie Extended model resulted in stronger magnitudes in the rostral cerebellum.

**Figure 2.**
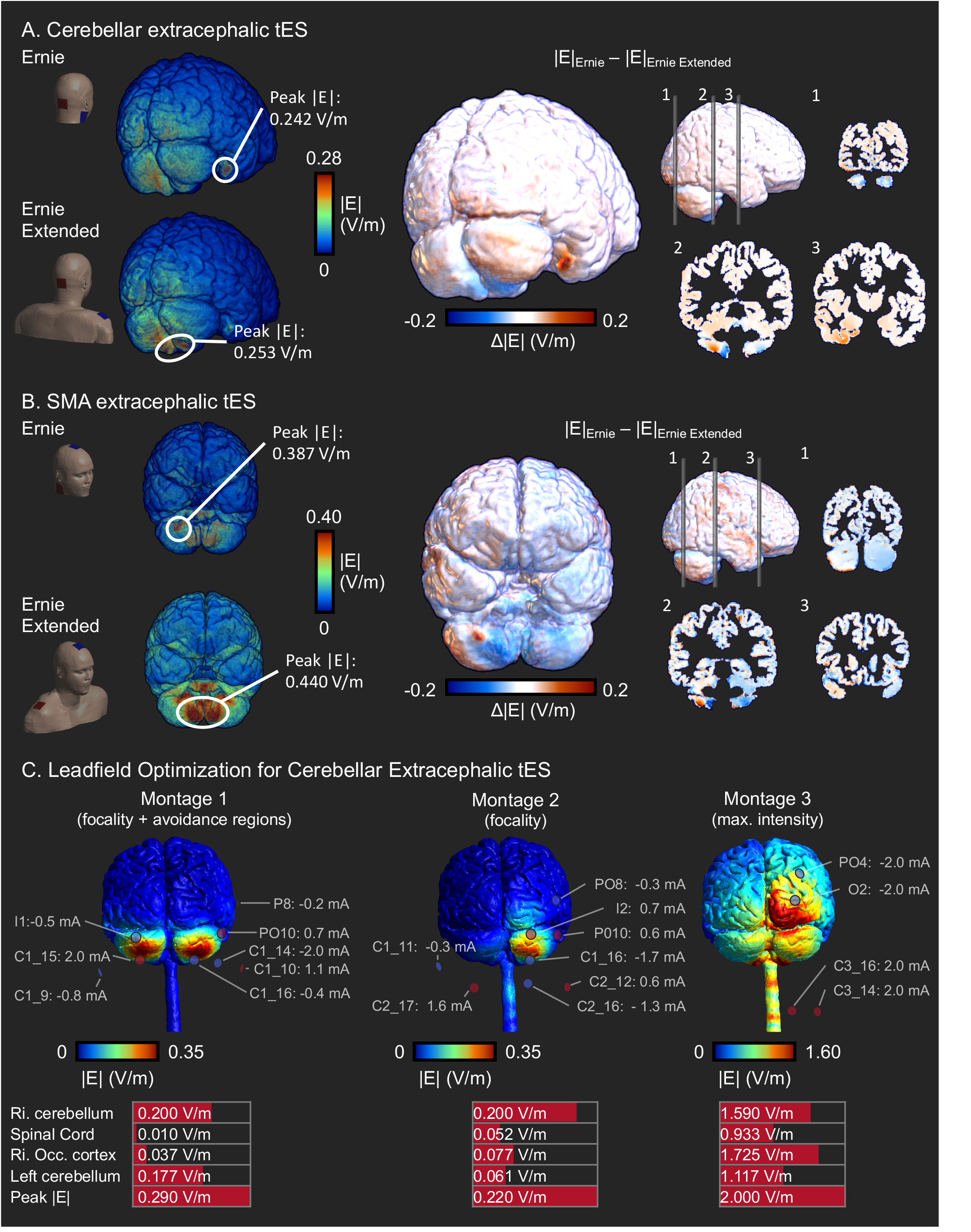
Differences in the modeled electric field between Ernie and Ernie Extended, and tES optimization. **A)** Electric field magnitude (|E|) difference between Ernie Extended and Ernie for cerebellar tES. While peak |E| was similar, its location differed. **B)** Difference for SMA extracephalic tES. Both peak |E| and peak location differed. **C)** Cerebellar tES optimization results in Ernie Extended. The upper figures show |E|, the lower table shows |E| in several regions of interest, and the peak magnitude. Montage 1 avoids co-stimulation of the spinal cord and right occipital cortex. Montages 2 and 3 aim for focality and intensity, respectively.

The extracephalic montage targeting the SMA induced larger magnitudes in the cerebellum than cerebellar tES, albeit having generally low focality. The peak magnitude of SMA tES in Ernie Extended was 0.440 V/m and 0.387 V/m in Ernie. The grey matter areas exceeding the 99.9th percentile magnitude again highlighted differences in the modeled electric fields: In Ernie Extended, peak fields were in the caudal cerebellum, while in Ernie, it was in the ventral cerebellum.

### 3.2. Effect of Extended Head Models on tES Optimization

We tested the utility of the extended head model that allows extracephalic electrode locations for optimizing multi-electrode tES targeting the right cerebellum (**Figure 1C**). We compared three montages as illustrated in **Figure 2C**. The value of Ernie Extended is highlighted by the inclusion of electrode locations outside of the standard head models in all montages.

Montage 1, aiming for focality while avoiding co-stimulation of the spinal cord and right occipital cortex, induced 0.200 V/m in the right cerebellar target. The peak induced electric field magnitude was 0.290 V/m and 9.64 cm^3^ of grey matter was exposed to electric fields above the 70^th^ percentile. While electric fields in the avoidance regions were generally low (> 0.037 V/m), the left cerebellum, which was no avoidance region, received more stimulation (0.177 V/m).

Montage 2, focused on focality, also induced 0.200 V/m in the target, peak electric field magnitude was 0.220 V/m, and 5.66 cm^3^ of the grey matter exceeded an electric field above the 70^th^ percentile. Although focality increased compared to montage 1, higher fields were present in the spinal cord and occipital cortex (**Figure 2C**).

Montage 3, focused on intensity, induced 1.590 V/m in the target, while the peak magnitude was 2.000 V/m. Focality was poor as 16.9 cm^3^ grey matter exceeded the 70^th^ percentile, with the occipital cortex receiving the highest doses (**Figure 2C**).

## 4. Discussion

We introduce Ernie Extended as an open-source model including the neck and shoulders offering high anatomical accuracy and highlight its value for simulations and optimizations of tES montages featuring extracephalic electrodes.

Previously, Callejón-Leblic and Miranda (2021) [11] compared a head model stopping at the ear and the lower end of the neck and found that the impact of head model extent depends on the electrode montage. The differences between the head models were largest for fronto-occipital tES where the full model results in more shunting through the lower head. Extending this work, we show that electrical field modeling of tES montages with extracephalic electrodes benefit from an extended head model. This enables to place the extracephalic electrodes at their actual positions rather than using approximated positions at the bottom of clipped models. We show that the location of the peak electrical field, an indicator for where tES enacts its primary effects, as well as voxelwise electrical field values differ when simulating tES of the right cerebellum or the SMA based on a non-extended or extended head model. For cerebellar tES, our modelling results concur with Ramersad et al. [31], who reported that extracephalic tES with a similar montage primarily targets the inferior cerebellum.

We also used Ernie Extended to optimize multi-electrode cerebellar tES via electrode positions absent in conventional head models, again demonstrating the value of extended head models. Overall, our results agree with Parazzini et al. [32], who used an extended head model to demonstrate the feasibility of focal cerebellar tES.

Our work has limitations. We used only one head model due to the time-intensive nature of manual segmentations, although anatomical idiosyncrasies can significantly affect electric field simulations [2]. Notably, previous work has demonstrated Ernie to be a reasonable group-average model [33]. Also, tissue conductivities are uncertain and strongly affect electric field simulations [34]. It remains to be demonstrated that distinguishing more tissue types indeed increases simulation accuracy. Lastly, our optimizations use eight electrodes, which are not always available. As our model and code are open-source, researchers can freely run optimizations for their available hardware.

## CRediT statement

Conceptualization: AT.

Methodology: SVH; VC; LDH; OP; AT.

Software: SVH; VC; LDH; OP; AT.

Validation: SVH; AT.

Formal analysis: SVH; AT.

Resources: HRS; RLJM; AT.

Data Curation: SVH; AT.

Writing - Original Draft: SVH; AT.

Writing - Review & Editing: SVH; VC; LDH; OP; HRS; RLJM; AT.

Visualization: SVH.

Supervision: HRS; RLJM; AT.

Project administration: HRS; RLJM; AT.

Funding acquisition: SVH; HRS; AT.

## Data Availability Statement

The Ernie Extended Model and accompanying code will be made freely available via the SimNIBS website.

## Appendices

### Appendix 1. Comparison of Ernie Extended with existing extended head models

| Model | Ernie Ex-<br>tended | *Callejón-*<br>*Leblic,*<br>*M.A. & Mi-*<br>*randa, P.C*<br>.[11] | *Im, C.-H., et*<br>*al., [12] Pa-*<br>*razzini et al.,*<br>*[16][32]* | *Datta, A., et al.,*<br>*[13]* | *Noet-*<br>*scher,*<br>*G.M., et*<br>*al.,*<br>*[14]* | *Mehta, A.R.,*<br>*et al., [15]* |
| --- | --- | --- | --- | --- | --- | --- |
| Open-source | ✓ | × | × | × | × | × |
| Underlying<br>data | T1w MRI<br>+ T2w<br>MRI + CT | ICBM152<br>v2009b &<br>ICBM152<br>v6 | Virtual<br>Family<br>Project | T1w MRI,<br>T2w MRI | Visible<br>Human<br>Project | T1w MRI |
| Lowest spi-<br>nal level | ~T5 | ~C7 | ~T11 | ~C4, syn-<br>thetic neck &<br>shoulders | ~T5 | ~T3, no<br>shoulders |
| Gyrification | ✓ | ✓ | × | ✓ | × | ✓ |
| Number of<br>tissues | 13 | 7 | 8 | 9 | 13 | 6 |

### Appendix 2. Scanning Parameters

Ernie Extended was created by means of four scans.

The high-resolution structural MR scans were acquired on a 3.0 T Philips Achieva MRI scanner using a 32-channel head coil. The T1w scan was acquired with the following parameters: Repetition time = 2600 ms; inversion time = 747 ms; echo time = 2.7 ms; flip angle = 8°; field of view =256×256× 208 mm^3^; voxel size = 1.0 mm^3^; band-width = 288.4 Hz; SENSE factor 2.5 along AP direction. The T2w scan was acquired with the following parameters: Repetition time = 2500 ms; echo time = 250 ms; flip angle = 90°; 224 sagittal slices; field of view = 245 × 245 × 190 mm^3^; voxel size =1.0 mm^3^; bandwidth = 969.2 Hz, SENSE factor 2 along AP and 1.8 along RL.

The T1w scan of the lower head and shoulders was acquired on the same scanner using the body coil for receive and the following settings: Repetition time = 2600 ms; inversion time = 534 ms; echo time =1.9 ms; flip angle = 8°; field of view = 300×528×350 mm^3^; voxel size = 2.0 mm^3^; bandwidth =378.1 Hz

The low-dose CT-scans was acquired on a Siemens Biograph mCT (PET-CT) scanner, with the following parameters: axial slices voxel sizes = 0.42 × 0.42 mm^2^; field of view = 215 × 215 mm^2^; resolution along Z-direction = 0.60 mm^3^; the z-direction extend was adjusted to cover the full neck while minimizing radiant dose; tube current-time produce = 115 mAs; tube potential = 80 KeV; maximum effective dose < 0.35 mSv.

### Appendix 3. Conductivity Values

| # | Tissue | Conductivity (S/m) | Reference |
| --- | --- | --- | --- |
| 1 | White Matter | 0.126 | [35] |
| 2 | Grey Matter | 0.275 | [35] |
| 3 | Cerebrospinal Fluid | 1.654 | [35] |
| 4 | Bone | 0.010 | [35] |
| 5 | Skin | 0.465 | [35] |
| 6 | Eye Balls | 0.500 | [3] |
| 7 | Compact Bone | 0.008 | [3] |
| 8 | Spongy Bone | 0.025 | [3] |
| 9 | Veins and arteries | 0.600 | [36] |
| 10 | Muscle | 0.160 | [36] |
| 11 | Cartilage | 0.880 | [37, 38] |
| 12 | Fat | 0.078 | [36] |
| 13 | Air | 0 |  |

